# Analgesia linked to Nav1.7 loss of function requires μ and δ opioid receptors

**DOI:** 10.1101/297184

**Authors:** Vanessa Pereira, Queensta Millet, Jose Aramburu, Cristina Lopez-Rodriguez, Claire Gaveriaux Ruff, John N Wood

**Affiliations:** Molecular Nociception Group, WIBR, UCL Gower Street London WC1E 6BT UK; Immunology Unit Department of Experimental and Health Sciences, Universitat Pompeu Fabra Carrer Doctor Aiguader No88, 08003 Barcelona, Spain; Institut de Génétique et de Biologie Moléculaire et Cellulaire, Université de Strasbourg, Centre National de la Recherche Scientifique, UMR7104, Institut National de la Santé et de la Recherche Médicale, U 1258, Ecole Supérieure de Biotechnologie de Strasbourg, Illkirch, France

**Author notes:** Correspondence to John Wood or Claire Gaveriaux-Ruff.

## Abstract

Functional deletion of the *SCN9A* gene encoding sodium channel Nav1.7 makes humans and mice pain-free (1,2). Opioid signaling contributes to this analgesic state (3). Here we show that the pharmacological block or deletion of both μ and δ opioid receptors is required to abolish Nav1.7 null opioid-related analgesia.κ-opioid receptor antagonists were without effect. Enkephalins encoded by the *Penk* gene are upregulated in Nav1.7 nulls (3). Deleting *Nfat5*, a transcription factor with binding motifs upstream of *Penk* (4), induces the same level of enkephalin mRNA expression as found in Nav1.7 nulls, but without consequent analgesia. These data confirm that a combination of events linked to SCN9A gene loss is required for analgesia. Higher levels of endogenous enkephalins (3), potentiated opioid receptors (5), diminished electrical excitability (6,7) and loss of neurotransmitter release (2,1) together contribute to the analgesic phenotype found in Nav1.7 null mouse and human mutants. These observations help explain the failure of Nav1.7 channel blockers alone to produce analgesia and suggest new routes for analgesic drug development.

---

Pain is numerically the greatest clinical challenge of the age, affecting about half the population, whilst 7% of people have debilitating pain conditions (9). Finding new analgesic targets and drugs has proved challenging. One approach has been to identify the genes involved in human monogenic loss of pain conditions (10). The association of human gain-of-function mutations in Nav1.7 with enhanced pain phenotypes, and the pain-free state linked to loss of Nav1.7 expression focused considerable attention on this voltage gated sodium channel as a potential analgesic drug target (11). Nav1.7 is found in damage-sensing peripheral sensory neurons, sympathetic neurons and CNS structures like the hypothalamus, as well as in non-neuronal locations such as the pancreas. Deletion in all sensory neurons and sympathetic neurons abolishes acute, inflammatory and neuropathic pain, although some pain disorders such as oxaliplatin-evoked cold allodynia are retained (2,12).

As human and mouse Nav1.7 null mutants are effectively pain-free, this channel should be an excellent analgesic drug target. However, channel blockers are very weak analgesics (11,13). This is likely due to the fact that partial channel block cannot recapitulate the many physiological effects of gene deletion. This explanation is supported by experiments that show that only 100% channel block with very high dose tetrodotoxin can recapitulate some effects of gene deletion (3). In null mutants neurotransmitter release is diminished, and synaptic integration is also diminished. In addition, the opioid peptide enkephalins are upregulated in the absence of Nav1.7, and opioid receptor signaling is potentiated. Both of these latter events may be linked to loss of sodium ingress through Nav1.7 (3).

Consistent with an opioid component of analgesia, the opioid antagonist naloxone substantially reverses Nav1.7 loss-associated analgesia (3). We wondered which opioid receptors were involved in this process. Here, using pharmacological studies and opioid receptor knockout mice, we show that both μ and δ opioid receptors contribute to Nav1.7 null mutant analgesia and deleting both receptors mimics the effects of naloxone on Nav1.7 null analgesia in mice. In addition, we show that elevating enkephalin mRNA levels to those found in Nav1.7 nulls is not alone sufficient to cause measurable analgesia.

We first examined the role of μ-opioid receptors (MOR) in Nav1.7 null-associated analgesia (Figure 1,A B). Nav1.7 null mutant mice show dramatic thermal analgesia. Global deletion of MOR on a Nav1.7 null background had a small effect on acute heat pain behaviour (Figure 1A). This effect did not match the effects of naloxone that substantially diminished analgesia (Figure 1B). Consistent with this, naloxone further diminished the analgesic phenotype of Nav1.7 / MOR double mutant mice, demonstrating that MORs alone do not account for the opioid-mediated component of Nav1.7 null associated analgesia (Figure1B).

**Figure 1:**
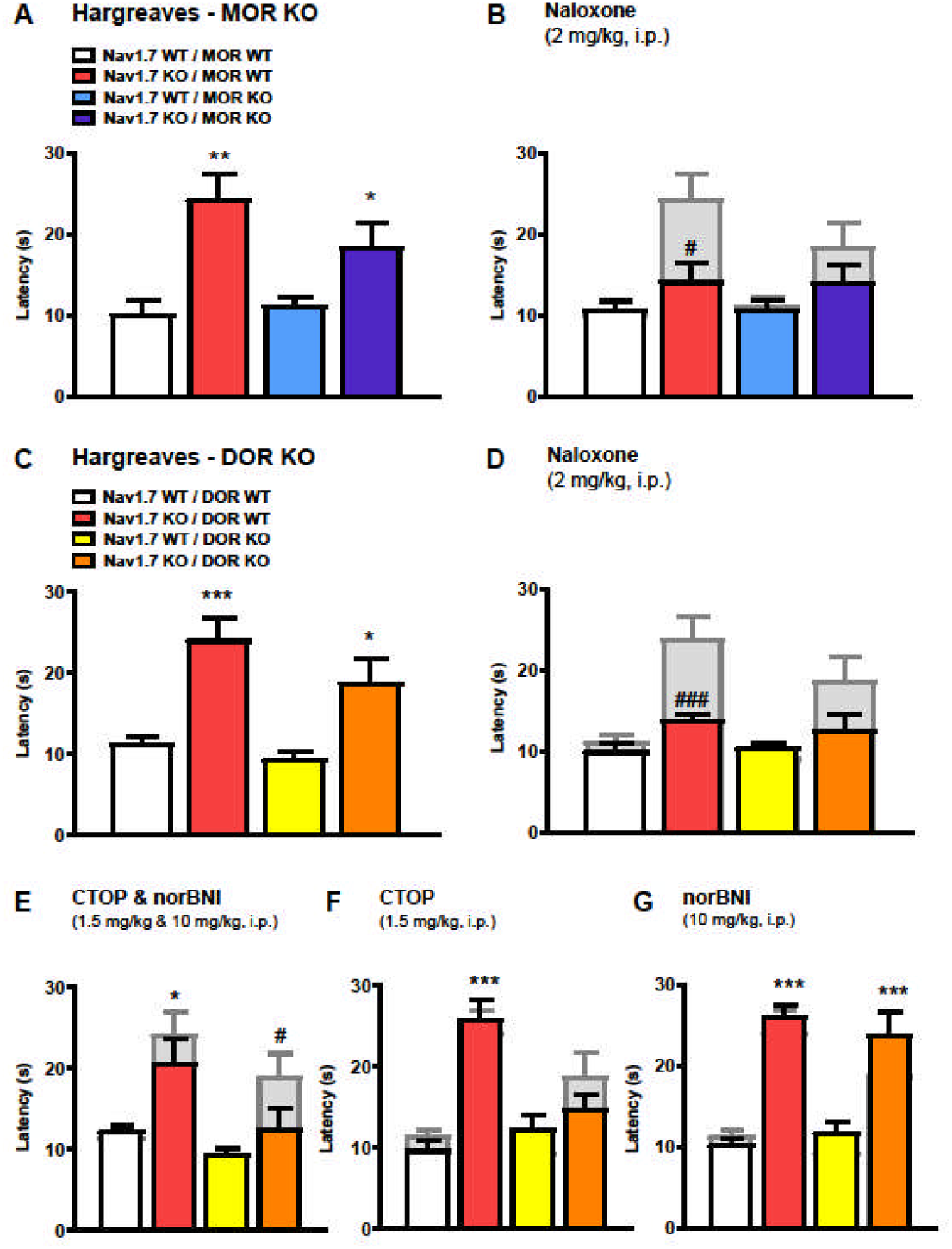
MOR or DOR deletion is not sufficient to reduce Nav1.7 KO pain sensitivity. (**A**) Noxious thermal stimulation of Nav1.7 WT/ MOR WT (white), Nav1.7 KO / MOR WT (red), Nav1.7 WT / MOR KO (blue) and Nav1.7 KO / MOR KO (purple) mouse hindpaw using Hargreave’s apparatus (n=8 per group). The withdrawal latency in seconds was manually recorded. (**B**) Hindpaw withdrawal latency 20 min after naloxone administration (2 mg/kg, i.p). The grey bars represent noxious thermal withdrawal latency baselines merged with latency measured 20 min after naloxone to facilitate comparison between pre and post drug injection pain-related behaviour. Results are presented as means ± SEM. Data were analysed by one-way ANOVA followed by Dunnett’s post hoc test (**A**) or two-way ANOVA followed by the Bonferroni post hoc test (**B**). * p<0.05 ** p<0.01 and *** p<0.001 vs Nav1.7 WT / MOR WT, & p<0.05 && p<0.01 and &&& p<0.001 vs own baseline). (**C**) Noxious thermal stimulation of Nav1.7 WT / DOR WT (white), Nav1.7 KO / DOR WT (red), Nav1.7 WT / DOR KO (yellow) and Nav1.7 KO / DOR KO (orange) mice (n=8 per group).(**D**) Hindpaw withdrawal latency 20 min after naloxone administration (2 mg/kg, i.p.). (**E**) Thermal withdrawal latency after a combination of the MOR antagonist CTOP (1.5 mg/kg, i.p.) and the kappa antagonist norBNI (10mg/kg, i.p.,) injected respectively 15 and 60 min before the test. (**F**) Effect of CTOP and (**G**) norBNI on mouse hindpaw withdrawal latency using Hargreave’s test (administrated 15 min or 60 min before recording the latency). Results are presented as mean ± SEM. Data were analysed by one-way ANOVA followed by Dunnett’s post hoc test (**C**) or two-way ANOVA followed by the Bonferroni post hoc test (**D**-**G**). * p<0.05 ** p<0.01 and *** p<0.001 vs Nav1.7 WT / DOR WT, & p<0.05 && p<0.01 and &&& p<0.001 vs own baseline.

Next we tested the effect of deleting δ-opioid receptors (DOR) on Nav1.7 null pain behaviour (16). Once again there was a small diminution in analgesia compared to Nav1.7 null mice (Figure1C). Naloxone further diminished the analgesic phenotype of the Nav1.7/DOR double null mutants (Figure 1D), demonstrating that DORs alone do not account for the opioid mediated component of Nav1.7 null associated analgesia. However, when the potent selective MOR antagonist CTOP was applied to DOR receptor null mice (17), the analgesia associated with Nav1.7 deletion was reduced by the same level as with naloxone (Figure 1F). CTOP and the kappa-opioid (KOR) antagonist norBNI (18) together also had the same effect as naloxone when applied to a Nav1.7/DOR double null (Figure1E). However, norBNI on a Nav1.7/DOR null background was without effect (Figure1G). These data show that KORs do not mediate analgesia in Nav1.7 null mutants, but pharmacological block of MOR on a DOR null background can account for all opioid mediated analgesia (17).

To provide further evidence that both MOR and DOR contribute to opioid-mediated analgesia in Nav1.7 nulls, we generated double opioid receptor null mutant mice on a Nav1.7 null background. Double MOR/DOR knockouts on a Nav1.7 null background showed exactly the same loss of analgesia as that caused by naloxone in Nav1.7 knockout mice (Figure2A,B). Application of μ δ and κ antagonists (19) together did the same (Figure2D), although the KOR antagonist norBNI alone was inactive, confirming that KOR activation did not contribute to analgesia (Figure2C). These pharmacological and genetic studies demonstrate that MOR and DOR together account for opioid-mediated analgesia in Nav.7 null mutant mice.

Elevated levels of enkephalins are found in Nav1.7 null mutant mice (3). Interestingly, there are five consensus binding sites for the transcription factor NFAT5 upstream of the *Penk* coding region. NFAT5 recognizes DNA elements similar to those bound by NFATc proteins (4). As NFAT5 activity is regulated by hyperosmolarity and salt kinases (20), there is a potential link between sodium ingress through Nav1.7 and transcriptional regulation. We manipulated sodium levels in sensory neuron cultures using either monensin as a sodium ionophore ([Na^+^] control 6.65 mM, SEM 0.27; [Na^+^] Monensin 9.46 mM, SEM 0.44; n = 19;) or veratridine as an activator of voltage gated sodium channels (control [Na^+^] 5.5 mM, SEM 0.25; [Na^+^] Veratridine 7.6 mM, SEM 0.41; n = 9) to increase sodium levels, and very high doses of TTX (500nM) to block voltage gated sodium channel activity and potentially lower intracellular sodium (Supplementary data). Intriguingly, agents that alter intracellular sodium concentrations impact similarly on Nav1.7 and NFAT5 mRNA levels. Monensin (Figure 3A) lowered both PENK and NFAT5 mRNA levels, whilst TTX elevated them (Figure3B). The TTX effect was apparent in wild type mice, but not in Nav1.7 nulls, implying that this channel is the locus of action for PENK mRNA control by TTX (Figure3C,D).

**Figure 2:**
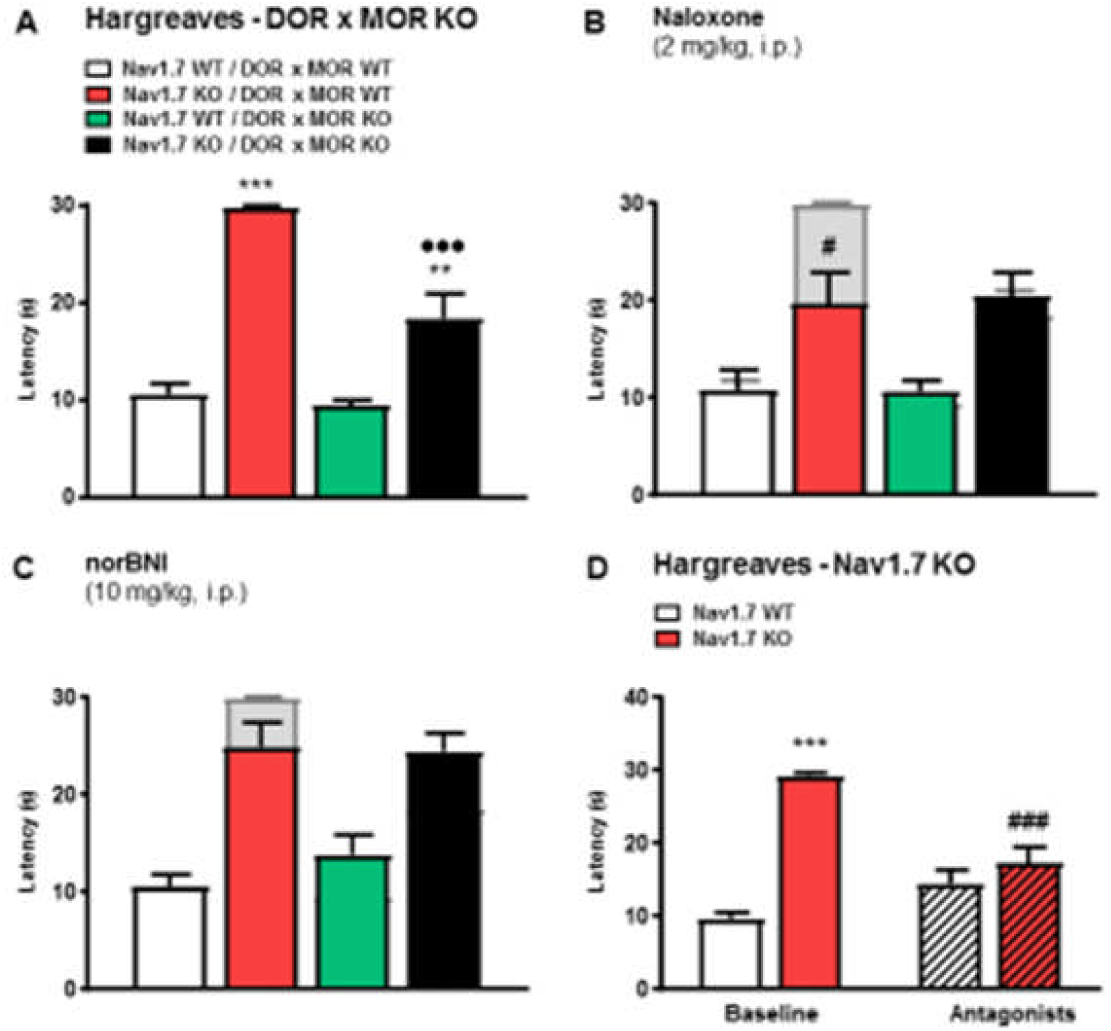
Deletion of both MOR and DOR mimics Naloxone effects on Nav1.7 KO pain thresholds. (**A**) Noxious thermal stimulation of Nav1.7 WT / DOR x MOR WT (white), Nav1.7 KO / DOR x MOR wt (red), Nav1.7 wWT/ DOR x MOR KO (green) and Nav1.7 KO / DOR x MOR KO (black) mice hindpaw using Hargreave’s apparatus (n=8 per group). (**B**) Hindpaw withdrawal latency 20 min after naloxone administration (2 mg/kg, i.p., saline). (**C**) Thermal withdrawal latency after administration of norBNI (10mg/kg, i.p.) injected 60 min before the test. Results are presented as mean ± SEM. Data were analysed by one-way ANOVA followed by Dunnett’s post hoc test (**A**) or two-way ANOVA followed by the Bonferroni post hoc test (**B**-**C**). * p<0.05 ** p<0.01 and *** p<0.001 vs Nav1.7 WT / DOR x MOR WT, # p<0.05 ## p<0.01 and ### p<0.001 vs own baseline. (**D**) Hindpaw withdrawal latency after administration of a combination of CTOP, NTI and norBNI (respectively 2 mg/kg, i.p., saline, injected 15 min before the test; 5 mg/kg, s.c., 30 min before and 10mg/kg, i.p. 60 min before the test) in WT (white bars) or Nav1.7 KO mice (red bars). Co injection of m d and k antagonists restores Nav1.7 KO thermal sensitivity. Results are presented as mean ± SEM. Data were analysed by two-way ANOVA followed by the Bonferroni post hoc test. * p<0.05 ** p<0.01 and *** p<0.001 vs Nav1.7 WT, # p<0.05 ## p<0.01 and ### p<0.001 vs baseline.

**Figure 3:**
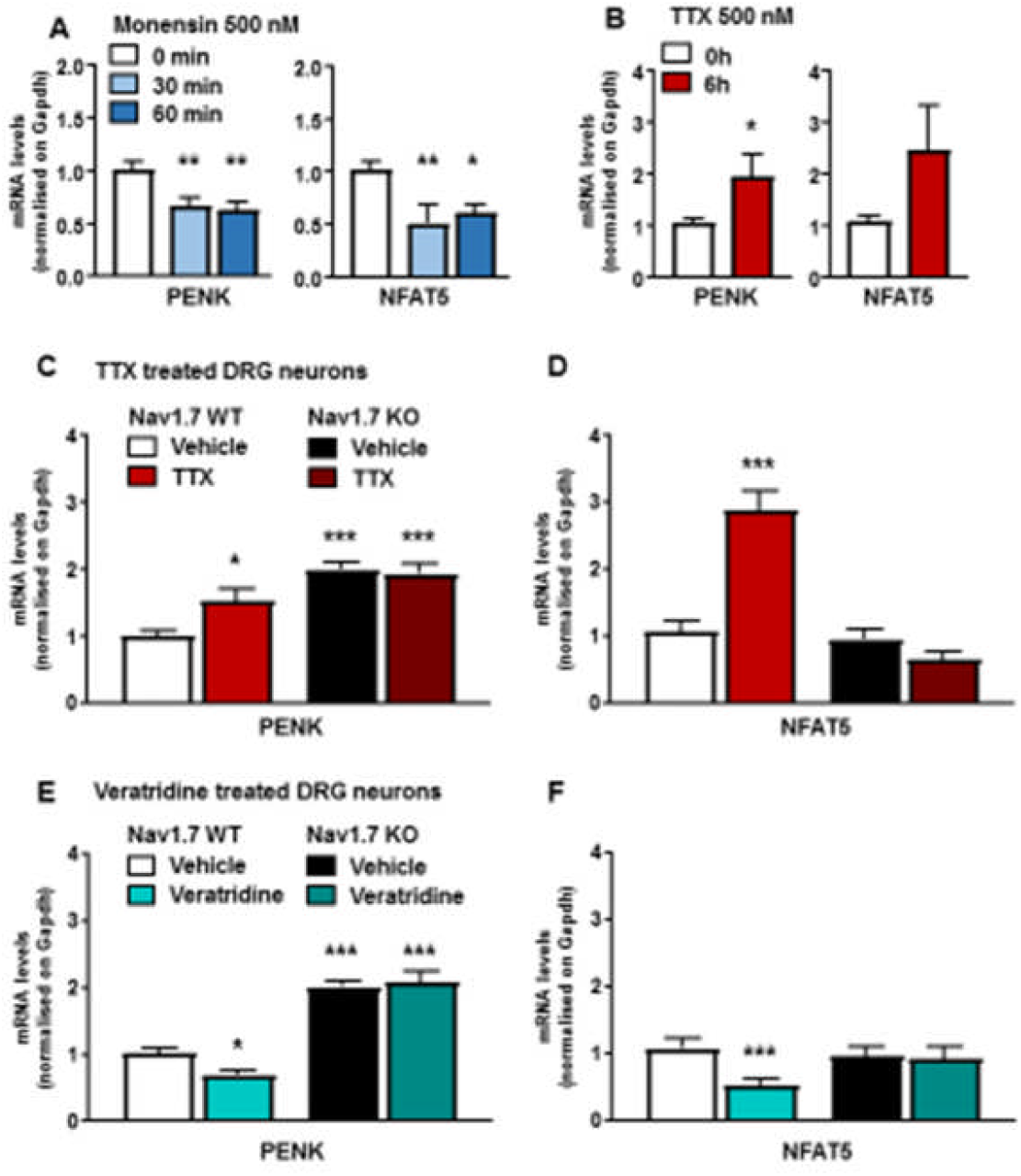
Both PENK and NFAT5 expression are regulated by intracellular sodium concentration. (**A**) Penk and Nfat5 expression levels in cultured DRG neurons treated with monensin (500 nM, 30 and 60 min, respectively grey and black bars). Control neurons (white bar) were treated with vehicle (ethanol) for 60 min. (**B**) Penk and Nfat5 mRNA quantification in cultured DRG neurons treated with TTX (500 nM, 6h). Control neurons received same volume of saline solution for 6h (black bar). (**C**) Penk and (**D**) Nfat5 transcripts levels in wt compared to Nav1.7 KO DRG neurons treated by TTX (500 nM, 6h). TTX induced Penk overexpression is correlated with Nfat5 expression level, both are dependant of Nav1.7 (**E**) Penk and (**F**) Nfat5 expression in wt and Nav1.7 KO cultured DRG neurons treated with veratridine (1 µM, 6h). Results are presented as mean ± SEM. Data were analysed by two-way ANOVA followed by the Bonferroni post hoc test. * p<0.05 ** p<0.01 and *** p<0.001 vs Nav1.7 WT Vehicle.

Veratridine lowered both PENK and NFAT5 mRNA levels in wild type but not Nav1.7 null mutant mice, again linking transcriptional events to Nav1.7 channel activity (Figure3E,F). We examined the role of NFAT5 using conditional NFAT5-Wnt1-Cre null mutants in sensory neurons of wild type and Nav1.7 null mutant mice. Expression levels of NFAT5 and Nav1.7 transcripts in single and double mutants were analysed to confirm Cre activity at the floxed loci (Supplementary data). NFAT5 conditional null mutant mice showed enhanced expression of PENK mRNA (Figure 4A). When the NFAT5 null mice were crossed with Nav1.7 null mutants, PENK mRNA levels further increased (Figure 4A). As NFAT5 null mice have the same levels of PENK mRNA as Nav1.7 null mutants, this allowed us to examine the contribution of enhanced opioid peptide expression to the analgesia seen in Nav1.7 null mutant mice. Opioid signalling in Nav1.7 null mutants is potentiated in at last two ways. First, there are enhanced levels of enkephalins, and second the opioid receptors have much enhanced activity, as measured indirectly through the quantitation of PKA signalling (5). There was, perhaps surprisingly, no analgesic effect of elevated enkephalin levels in the *Nfat5* null sensory ganglia. By measuring noxious mechanosensation (Figure 4B), thermal thresholds (Figure 4C), and noxious heat-induced pain related behaviour (Figure 4D), the NFAT5 null enkephalin-induced mice showed normal pain behaviour, compared to Nav1.7 null mice (Figure 4). As opioids clearly play a role in Nav1.7 null analgesia, as demonstrated by the naloxone effects, this suggests that the enhanced activity of opioid receptors may make a major contribution to the Nav1.7 null opioid-mediated analgesia.

**Figure 4:**
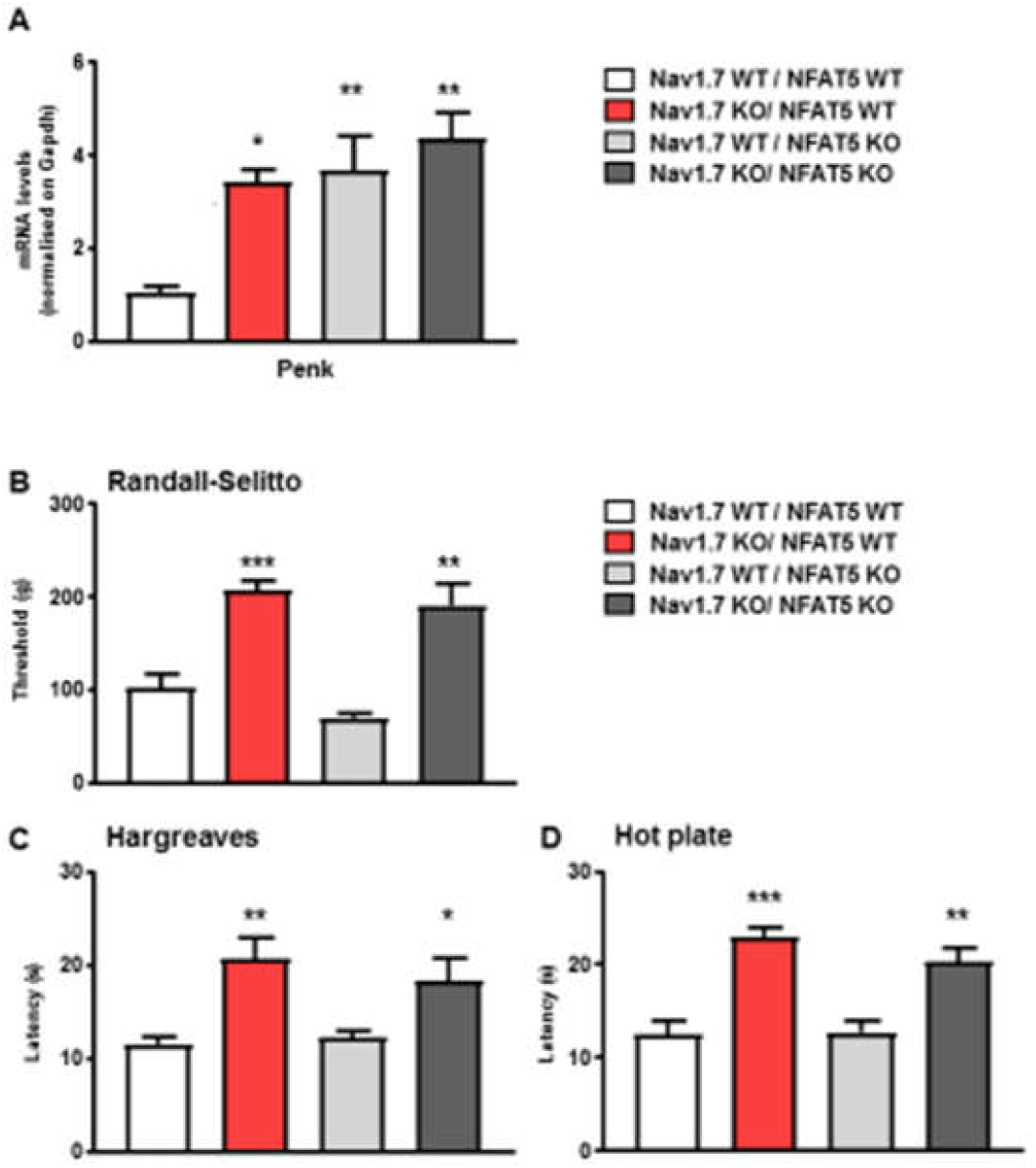
*Nfat5* conditional gene deletion induces PENK overexpression *in vivo* without eliciting a pain-insensitive phenotype. (**A**) Expression levels of PENK transcript in Nav1.7 WT/ NFAT5 WT (white bar), Nav1.7 KO / NFAT5 WT(red bar), Nav1.7 WT / NFAT5 KO (light grey) and Nav1.7 KO / NFAT5 KO mice (dark grey). (**B**) Noxious mechanical pressure threshold of the same four mice lines using the Randall-Selitto apparatus. (**C**) Noxious thermal stimulation of Nav1.7 WT/ NFAT5 WT (white bar), Nav1.7 KO / NFAT5 WT(red bar), Nav1.7 wt / NFAT5 KO (light grey) and Nav1.7 KO / NFAT5 KO mice (dark grey) mice hindpaw using Hargreave’s apparatus (n=8 per groups). (**D**) Response to noxious thermal stimulation by using the hotplate test at 55°C. Results are presented as mean ± SEM. Data were analysed by one-way ANOVA followed by the Dunnett’s post hoc test. * p<0.05 ** p<0.01 and *** p<0.001 vs Nav1.7 WT / NFAT5 wt.

What are the implications of these findings for drug development? Firstly the complexity of physiological changes that occur in Nav1.7 nulls is striking. Receptors (e.g. 5HTr4) and transcription factors (e.g. Runx1) implicated in nociception are dysregulated (3) Opioid peptide expression is increased (3) and opioid signalling is potentiated (5), whilst electrical excitability (6) and integration of nociceptive stimuli is lost (7). There is evidence that these events require the complete loss of Nav1.7 function, as occurs in null mutants. For example, only complete channel block with very high doses of TTX can induce increased PENK mRNA expression (3). Should small molecule-specific Nav1.7 antagonists be able to replicate all these events then they would be excellent analgesics. All the evidence thus far demonstrates that this is not the case, and the necessarily partial blockade of Nav1.7 does not cause analgesia (13). Molecules with limited specificity like Biogen’s BIIB074 are good analgesics but much of their activity likely results from blockade of sodium channels other than Nav1.7 (20).

The role of MOR and DOR and the lack of a role for KOR in Nav1.7 null analgesia fit with recent data. There is evidence for μ-δ receptor interactions in nociceptive sensory neurons (21), and primates express μ-δ heteromultimers as targets of opioid analgesia (22). As Nav1.7 deletion with peripheral nervous system-dependent Cre mice causes analgesia, then the actions on opioid receptors must occur either on primary sensory neurons, or on their synaptic targets within the spinal cord. Evidence that co-administration of opioids with Nav1.7 antagonists can have synergistic therapeutic effects has been demonstrated with a number of specific Nav1.7 antagonists. However, human proof of concept studies on synergistic analgesia with Nav1.7 antagonists and opioids have yet to be published. The evidence for potentiation of opioid receptor signalling in Nav1.7 null mice is significant (5). Although diminished electrical excitability may provide the necessary landscape for endogenous opioid effects, it is surprising that elevated enkephalin levels alone do not produce any detectable levels of analgesia in the NFAT5 null mice. Exogenous administration of enkephalins in humans delivered through gene therapy has useful analgesic effects (23). The focus is then upon potentiated opioid receptor signaling (5). There is some evidence linking the ingress of sodium through Nav1.7 to effects on GPCR activity. Pert and Snyder showed the influence of sodium on opioid receptor activity in 1974, demonstrating that increased sodium concentrations caused diminished agonist binding (*24*). Intracellular sodium levels may control this process (25) and the proximity of Nav1.7 channels to opioid receptors may influence sodium occupancy of these GPCRs (26).

In summary, μ and δ opioid receptors are required for the opioid component of Nav1.7 null mutant analgesia. Co administration of μ/δ agonists with specific Nav1.7 antagonists may therefore have useful analgesic effects (29). If analgesia depends substantially upon both potentiated receptor activity, as well as increased enkephalin expression, analgesic drug development using small molecules to mimic Nav1.7 gene deletion will be problematic. Nociceptor silencing through CRISPR mediated gene deletion of Nav1.7 may prove a more tractable analgesic strategy for extreme chronic pain conditions (30).

## Acknowledgements

We thank the Wellcome Trust (grants **101054/Z/13/Z, 200183/Z/15/Z**) and Arthritis Research UK (20200) for generous support. We thank David Reiss at IGBMC for providing the colonies of opioid receptor mutant mice. We thank James Cox, Jing Zhao and members of the Molecular Nociception Group for comments and advice.

## Material & Methods

### Animals

Nav1.7 floxed mice were generated as described (27). Specific deletion of SCN9A exons 14 and 15 was performed by crossing Nav1.7^flox/flox^ mice with Wnt1-Cre^tg/0^ hemizygous transgenic mice purchased from Jackson Labs (129S4.Cg-Tg(Wnt1-cre)2Sor/J, Stock No: 022137). F1 offspring was crossed to obtain Nav1.7^flox/flox^:Wnt1-Cre^tg/0^ and further bred with either MOR^−/−^ or DOR^−/−^ mice. Previously reported MOR and DOR null mutants (Filliol et al., 2000; Matthes et al., 1996) were used. We obtained Nav1.7^flox/flox^:MOR^−/−^:Wnt1-Cre^tg/0^ and Nav1.7^flox/flox^:DOR^−/−^:Wnt1-Cre^tg/0.^ Finally, triple mutants carrying either MOR or DOR homozygous deletion were crossed in order to generate Nav1.7^flox/flox^:MOR^−/−^:DOR^−/−^:Wnt1-Cre^tg/0^. For all mouse lines, homozygous mutants were compared to Wnt1-Cre negative animals. For clarity, Nav1.7^flox/flox^:DOR^+/+^:Wnt1-Cre^0/0^ are named in this article Nav1.7 WT / DOR WT; Nav1.7^flox/flox^:DOR^+/+^:Wnt1-Cre^tg/0^, Nav1.7 KO / DOR WT; Nav1.7^flox/flox^:DOR^−/−^:Wnt1-Cre^0/0^, Nav1.7 wt / DOR KO; and finally Nav1.7^flox/flox^:DOR^−/−^:Wnt1-Cre^tg/0^, Nav1.7 KO / DOR KO. The same simplification was applied for all the genotypes. NFAT5 floxed mice were generated by Dr Cristina López-Rodriguez (Barcelona, Spain). Experiments were conducted using both male and female that were between 8 and 12 weeks old at the time of experiments. Animals were housed up to 5 per cage, in a temperature-controlled room with a 12-h light–dark cycle. Food and water were available ad libitum. Genotyping was carried out on genomic DNA extracted from ear notches and PCR was conducted as described (16,17,27).

### Behavioural Testing

Animal experiments were approved by the UK Home Office and UCL ethics committee Act 1986. Mice were acclimatized to the experimental room and were handled during a period of 1 week before starting the experiments. Observers who performed behavioural experiments were blind to the genotype. *Hargreaves thermal test*: The animal’s hindpaw was exposed to an intense light beam and the withdrawal latency recorded. *Randall Selitto test*: A blunt probe was used to apply force approximately midway along the tail. *Hot plate test*: Animals were exposed to a 55°C chamber floor and the withdrawal latency recorded.

### Drugs

Naloxone, Naltrindole hydrochloride (NTI), CTOP and nor-Binaltorphimine dihydrochloride (norBNI) were purchased from Sigma, UK and dissolved in saline. They were respectively administered 30 min, 30 min, 15 min and 60 min before performing behavioural experiments. Unless specified, all drugs were injected intraperitoneally at the dose described in the figure legend (typically, 2 mg/kg for naloxone, 5 mg/kg for NTI, 1.5 mg/kg for CTOP and 10 mg/kg for norBNI). Monensin, TTX and Veratridine (Sigma, UK) were respectively dissolved in ethanol, saline and DMSO and incubated for the time specified in the figure legend. For controls, the same volumes of vehicle were used.

### DRG neuron cultures

DRG from all spinal levels were harvested and dissociated as described (28). Dissociated neurons were plated on poly-L-lysine and laminin coated 35-mm plastic dishes (Nunc, Denmark). Incubation with drugs was started at least 24h after dissociation. Monensin (Sigma, UK, in ethanol absolute), TTX (Sigma, UK, in extracellular solution) or Veratridine (Sigma, UK, DMSO) were used at concentrations described in the figure legends before RNA extraction and quantification. For each experiment, control DRG neurons were treated with the appropriate vehicle.

### Quantitative PCR

For fresh DRG analysis, DRG from lumbar segments L4, L5 and L6 were harvested and pooled. For DRG cultures, cells were collected after incubation with the drug and concentrated by centrifugation. RNA was extracted using TRIzol^®^ Reagent (Invitrogen) according to the manufacturer’s instructions. Reverse transcription was performed using iScript^™^ Reverse Transcription Supermix for RT-qPCR following Bio-Rad supplied protocol. cDNA amplification was performed in triplicate, using SsoAdvanced^™^ Universal SYBR^®^ Green Supermix (Bio-Rad). DNA amplification was quantified with a Bio-Rad CFX Connect^™^ Real-Time PCR Detection System thermocycler. The expression level of target genes was normalized to housekeeping gene mRNA (*Gapdh*). Fold changes were determined using the 2^−ΔΔCt^ equation in which wild-type littermate or vehicle-treated cultured DRG cDNA samples were designated as the calibrator. The data presented are given as the mean of the fold changes.

### Statistical analysis

Data were analysed using GraphPad Prism 7. Statistical tests performed for a given experiment are described in figure legends.

### Supplementary methods

Live cell imaging. For Na^+^ imaging, neurons were loaded for 30 min with 5 µM of SBFI in serum free DMEM, and then washed with extracellular solution (140 mM NaCl, 3 mM KCl, 10 mM HEPES, 10 mM D-Glucose, 2 mM CaCl2, 1 mM MgCl2, pH 7.4 adjusted with KOH, Osmolarity 300 mOsm adjusted with D-Glucose). Cells were alternately excited at 340 and 380 nm and emissions at 510 nm collected separately to determine 340/380 nm ratio. Calibration of [Na^+^]i was performed by exposing SBFI-loaded DRG neurons to different extracellular solutions with specific Na^+^ concentration for 30 min (in the additional presence of 3µM gramicidin D for equilibrium between intracellular and extracellular Na+ concentration). For Ca^2+^ imaging, cells were loaded with 1 µM of Fura-2 for 30 min and alternatively excited at 340 and 380 nm. Results were expressed using the ratio of the 340 nm/380 nm wavelengths.

